# Representing Transcription Factor Dimer Binding Sites Using Forked-Position Weight Matrices and Forked-Sequence Logos

**DOI:** 10.1101/2024.07.16.603695

**Authors:** Matthew Dyer, Roberto Tirado-Magallanes, Aida Ghayour-Khiavi, Xiao Xuan Lin Quy, Walter Santana, Hamid Usefi, Morgane Thomas-Chollier, Sudhakar Jha, Denis Thieffry, Touati Benoukraf

**Author notes:** Correspondence to: **Touati BENOUKRAF**, *PhD*, Division of BioMedical Sciences, Faculty of Medicine Craig L. Dobbin Genetics Research Centre, Room 5M318, Memorial University of Newfoundland St. John’s, NL A1B 3V6, Canada, **Denis Thieffry**, *PhD*, U900 Institut Curie, INSERM, Mines, Université PSL 26 Rue d’Ulm, 75005 Paris, France. These authors contributed equally to this work (Matthew Dyer, Roberto Tirado-Magallanes, Aida Ghayour-Khiavi). Bacterial infection, response & dynamics, Institut de Biologie de l’ENS (IBENS), Département de biologie, École normale supérieure, CNRS, INSERM, Université PSL, 75005 Paris, France.

## Abstract

Current position weight matrices and sequence logos may not be sufficient for accurately modeling transcription factor binding sites recognized by a mixture of homodimer and heterodimer complexes. To address this issue, we developed *forkedTF*, an R-library that allows the creation of Forked-Position Weight Matrices (FPWM) and Forked-Sequence Logos (F-Logos), which better capture the heterogeneity of TF binding affinities based on interactions and dimerization with other TFs. Furthermore, we have enhanced the standard PWM format by incorporating additional information on co-factor binding and DNA methylation. Precomputed FPWM and F-Logos are made available in the *MethMotif 2024* database, thereby providing ready-to-use resources for analyzing TF binding dynamics. Finally, *forkedTF* is designed to support the TRANSFAC format, which is compatible with most third-party bioinformatics tools that utilize PWMs. The *forkedTF* R-library is open source and can be accessed on GitHub at https://github.com/benoukraflab/forkedTF.

## INTRODUCTION

Position weight matrices (PWM) (1) and sequence logos (2) have become standard tools to model and visualize transcription factor binding sites (TFBS) on the aggregation of DNA sequences that are known to be recognized by a given transcription factor (TF). Initially, DNA binding sequences were characterized through in vitro studies such as gel-shift assays and, later, SELEX, which outputs up to 20 DNA sequence variants bound by a specific TF. Advances in high-throughput technologies have enabled the genome-scale characterization of TFBS sequences in an *in vivo* context, particularly through the use of Chromatin Immunoprecipitation of a TF of interest, followed by deep sequencing of the bound DNA loci (ChIP-Seq). ChIP-seq experiments and their derivates (3) aim to characterize TF binding in the natural cellular context, including chromatin modifications and the occurrence of transcription co-factors.

It is widely recognized that many TFs bind to DNA as dimers, such as those containing Leucine Zippers. Depending on the cellular context, these complexes can be composed of either two similar proteins (homodimer) or two distinct proteins (heterodimer). Therefore, ChIP-seq assays capture the entire collection of dimer combinations present in the cell. However, aggregating the heterogeneous DNA-bound sequences into a single PWM or sequence logo results in a bipartite or dyad motif consisting of a conserved part (binding sites of the TF of interest) and a degenerated part (aggregation of the binding sites of the main TF partners). As a result, current PWMs and sequence logos poorly model TF dimer binding affinities since the corresponding PWM segments are often noisy. To address this issue, we developed *forkedTF*, an R-library that enables the generation of Forked-Position Weight Matrices (FPMW) and Forked-Sequence Logos (F-Logos), which better depict the sequence affinity of a TF of interest along with those of a segregated list of partners.

## MATERIALS AND METHODS

### Overview of *forkedTF* workflow

The *forkedTF* package consists of four main functions, which can be used sequentially: *miniCofactorReport*, *createFPWM*, *plotFPWM* and *write.FPWM*. The first step aims to identify the main co-binding partners of a transcription factor in a particular cell line using the function *miniCofactorReport*, a TFregulomeR API (4) that extracts the relevant ChIP-seq and DNA methylation data from the *MethMotif* (5, 6) and *GTRD* databases (7). The resulting report contains a list of co-binding partners of the TF of interest, along with detailed information on TF co-localization, motif usage, and DNA-binding motif methylation profile. Using this report, users can select the optimal fork position to generate a Forked-Position Weight Matrix (FPWM) object using the *createFPWM* function. This object can be visualized as a Forked-Sequence Logo (F-Logo) using the *plotFPWM* function and exported as a TRANSFAC-compatible format file using the *write.FPWM* function (Figure 1).

**Figure 1.**
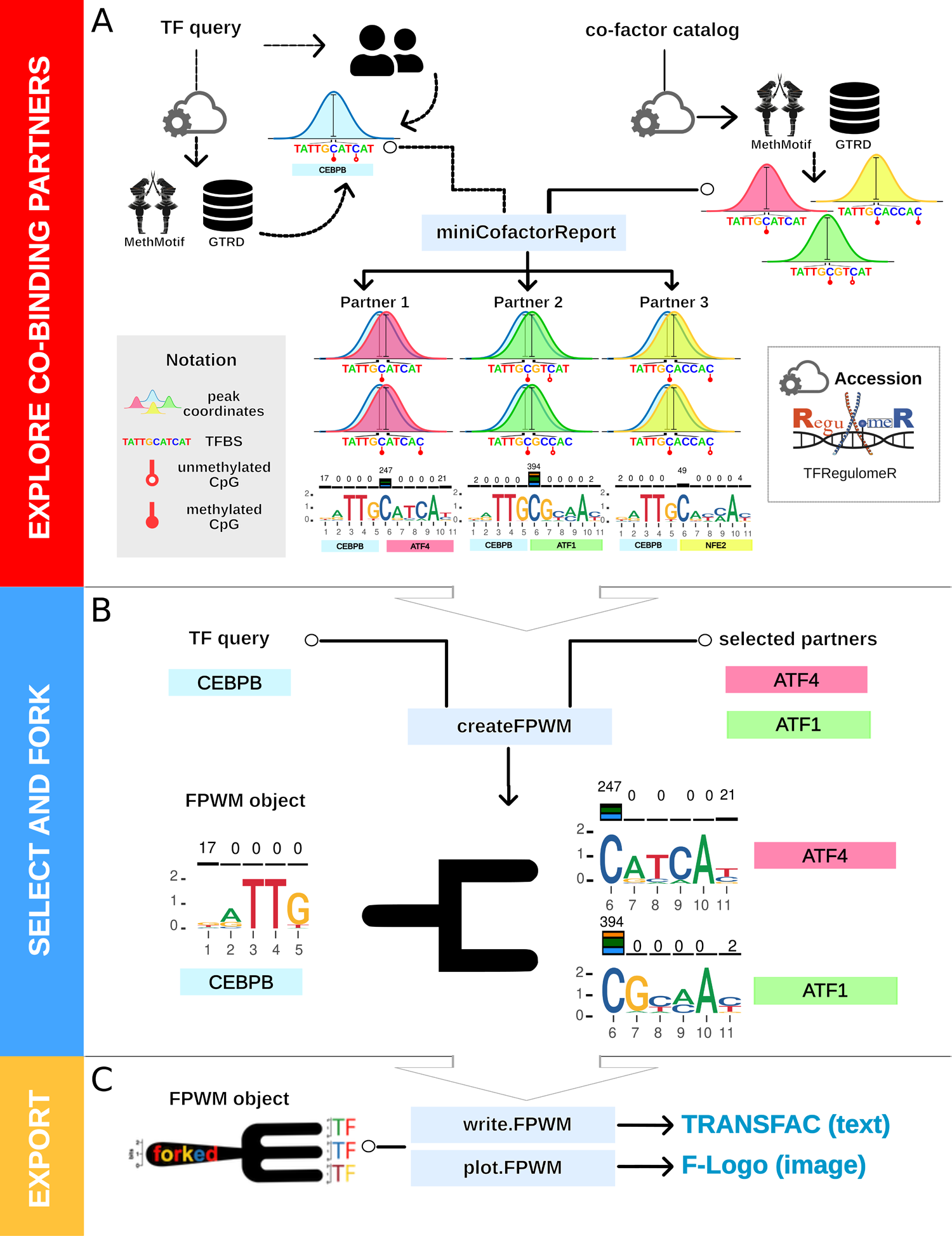
forkedTF workflow for generating FPWM objects, FPWM files, and F-Logo plots. **(A)** The initial step involves identifying the main binding partners of a transcription factor of interest using the *miniCofactorReport* function. This function utilizes a large cell-specific dataset compendium from the *MethMotif* and *GTRD* databases, with the option to integrate custom ChIP-seq peak lists in bed format. The output report provides information on the co-binding proportion, dimer motif usage, and DNA methylation profile for each pair of TFs. **(B)** This report enables the user to identify the portion of the motif corresponding to the main TF, which is crucial for setting the fork position. The *createFPWM* function then generates a Forked-Position Weight Matrix (FPWM) object. **(C)** This object can be exported in a standard TRANSFAC-compatible format file or visualized as an F-Logo plot using the *write.FPWM* and *plot.FPWM* functions, respectively.

### Step 1: miniCofactorReport()

The function *miniCofactorReport* interrogates the TFregulomeR data compendium to retrieve ChIP-seq and DNA methylation records in a cell-specific manner. As input, *miniCofactorReport* requires the ID of a TF of interest and a cell type. Then, the function queries all available ChIP-seq datasets within a specific cell type to identify the TF of interest’s partners. The function *GenomicRanges* (8) is then used to compute the intersection of ChIP-seq peak signals for each pair of TFs to detect co-binding. All PWMs within ChIP-seq peak intersections (+/- 100 bp region surrounding the main TFBS) are computed using MEME-ChIP (9). Next, DNA methylation is profiled for these intersected DNA loci and mapped to the binding motif. Each pair of TFs are then ranked by co-binding frequency, which is defined as the fraction of the number of loci bound by the reference TF that are co-bound by the cofactor. The user can provide a co-binding threshold for the report, which is set to 5% by default. Additionally, the over-representation of co-binding events is assessed by adjusted p-values computed using the *extractEnrichment* function from the ReMap package (https://github.com/remap-cisreg/ReMapEnrich).

Given the nature of the analysis, which involves large datasets of ChIP-seq and DNA methylation, users have the option to run the query online (on the Digital Research Alliance of Canada) or locally with *TFregulomeR* (4).

### Step 2: createFPWM()

The *createFPWM* function generates an FPWM object. It requires the ID of the TF of interest, the cell type of interest, and a list of co-factors obtained by *miniCofactorReport*, along with the chosen fork position, as inputs. The FPWM object is actually a list of PWMs for each TF of interest co-factor.

### Step 3: plotFPWM()

FPWM objects can be visualized using the *plotFPWM* function, which generates a PDF vector image containing a forked logo. In this logo, positions preceding the fork correspond to the core motif of the reference TF, which is connected *via* a fork with sequence logos corresponding to the binding partners. The sequence logos show the methylation levels for each cytosine in a CpG context located within the binding site. The proportion of binding overlap between the TF of interest and each binding partner is displayed on top of each forked arrow as a percentage.

### Step 4: writeFPWM()

FPWM objects can be stored as a series of TRANSFAC files or a TRANSFAC-like file (*i.e.* FPWMtransfac - Supplementary Figure 1) for further analysis using the function *writeFPWM*. In our suggested FPWMtransfac format, the metadata section contains several variables, including the ID of the main TF labelled “parentLogo” and the IDs of the partner TFs labelled “leafLogos.” Information regarding the overlap percentage, the number of base pairs in the overlap, and the total number of overlapping peaks are recorded in the “overlappingScore”, “numberOfBasePairs”, and “numberOfOverlappingPeaks” fields, respectively. Each comma-separated value in those fields corresponds to the TF in the same order as appeared in leafLogos (i.e., the first value in the “overlappingScore” field corresponds to the first TF in leafLogos, the second value to the second TF, and continuing to the nth value). The comma-separated value fields follow the same pattern for the nucleotide frequency matrix. The positions preceding the fork position correspond to the reference TF shown in the “parentLogo” field, whereas the values following the fork position correspond to the binding partners ordered as listed in the “leafLogos” field (Figure 2). Of note, because standard TRANSFAC matrices are derived from a set of DNA sequences of the same length, the sum of all the nucleotide counts remains the same for every row.

**Figure 2.**
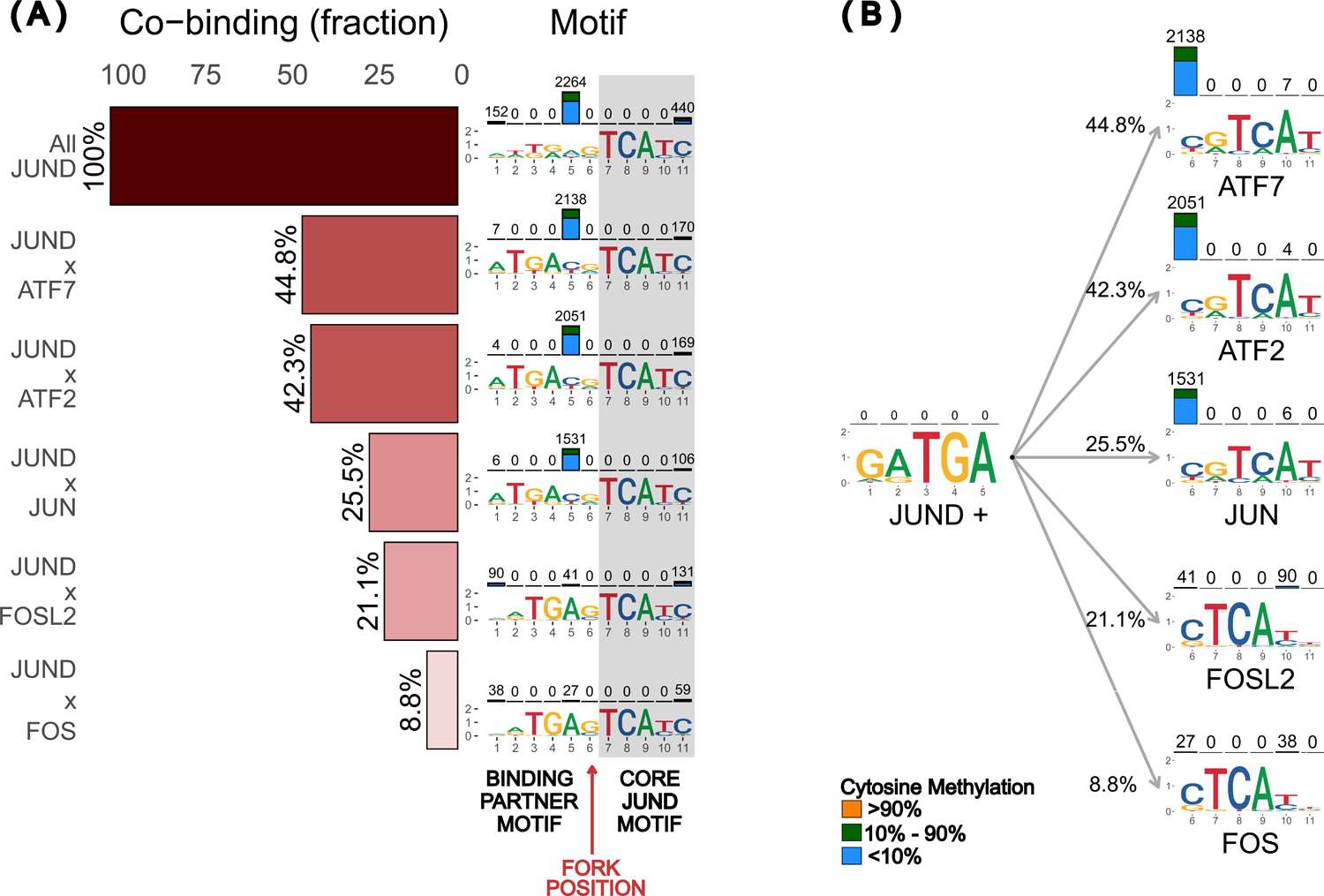
forkedTF analysis of JUND binding profile in HepG2 cell line. **(A)** *miniCofactorReport* uses the TFregulomeR data compendium to find the main binding partners of a TF of interest in a given cell type. Here, we show the output for JUND as the main factor and its co-binding partners in HepG2. The top six co-factors are shown, along with their percentage of peak overlap with JUND peaks (bar plot on the left) and a motif logo calculated from the peak overlap (right) with the cytosine DNA methylation on top (bars on top of the motif). The last five positions in the motif are conserved across all binding partners (highlighted), suggesting that this is the main TF’s core motif (JUND), while the rest of the position corresponds to the motif of the binding partner. **(B)** A forkedLogo representation of the *miniCofactorReport* results can be generated to illustrate the core motif of the main TF (left) forking into six co-factor motif logos (right).

However, this is not the case for FPWMs, where binding sequences are split between the main TFBS and TFBS belonging to co-factors. This specificity of FPWM may lead to some incompatibility with current scanning PWM algorithms. To overcome this potential issue, we have implemented three different matrix formats: (i) Count matrix, which records raw counts from the intersection of peaks; (ii) Probability matrix, where counts in rows are divided by the sum of the elements in these rows (i.e., the sum of elements in rows always equals 1); and (iii) Scaled Count matrix, where counts are scaled with respect to the highest value among the sum of elements for each row (Supplementary Figure 2).

### Comparative motif enrichment analysis

To perform the computations at the basis of panels B to F of Figure 4, we used several tools from the RSAT suite (10) on the web server https://rsat.france-bioinformatique.fr/metazoa/:

1. We have used *fetch-sequences* to extract 101bp-long sequences flanking the centers of the peaks resulting from the intersection of the JUND ChIP-seq peak set with the ATF2 (JUNDxATF2) or FOSL2 (JUNDxFOSL2) peak sets, in the HEPG2 cell line.
2. We have run *matrix-quality* (11) to compute the best scores of the three reference count matrices (previously generated with MEME-ChIP using all JUND peaks, JUNDxATF2 peaks, and JUNDxFOSL2 peaks, respectively), on the HEPG2 JUNDxATF2 and JUNDxFOSL2 peak sets, which were gathered to generate the score distributions shown in Figure 4B. We ran this analysis with a background model consisting of a Markov model of order two computed on GRCh37 Homo sapiens upstream non-coding sequences, together with pseudo frequencies set to 0.01. The matrix columns were randomly permuted five times to generate the negative control curve. The theoretical curve was computed directly from the background Markov model.
3. We have used *matrix-scan* (https://rsat.france-bioinformatique.fr/metazoa/matrix-scan_form.cgi) to scan the JUNDxATF2 and JUNDxFOSL2 peak sequence sets with the same three reference count matrices, with the option quick, a p-value threshold set to 0.01, a background Markov model of order 1, and the pseudo frequency parameter set to 0.01. Of note, as a convention, we use ‘x’ to represent intersections of peaks and ‘+’ to represent intersections of motifs, as illustrated below:

- JUNDxATF2 (or JUNDxFOSL2) is the intersection between the JUND peak set and the ATF2 (or FOSL2) peak set.
- JUND+ATF2 (or JUND+FOSL2) represents a specific (bi-partite) matrix or motif discovered in the set of peaks obtained by computing the intersection between the JUND peak set and the ATF2 (or FOSL2) peak set using MEME-ChIP.

## RESULTS

### Application of *forkedTF* to JUND binding in HepG2 cells

As an illustration of improvements made by *forkedTF*, we have performed an analysis of the transcription factor JUND in the HepG2 cell line. First, *MiniCofactorReport* was used to identify the following five co-factors of JUND: ATF7, ATF2, JUN, FOSL2 and FOS (Figure 2A, Supplementary Figure 3). In HepG2, we observe that the JUND global binding motif is highly conserved within the five last base pairs; however, the preceding bases (positions 1 to 6) are highly degenerated. Interestingly, the MiniCofactorReport results show that none of the JUND dimerized motifs present the observed degeneration before position seven. In addition, it can be noticed that the nucleotides in positions 7 to 11 are highly consistent across all binding partners, suggesting that these final five base pairs correspond to the JUND core motif.

In contrast, the variable motif sequences observed before position seven support the fact that they belong to distinct binding partners and can be categorized into two main groups. The first group is characterized by a double spacer in positions 5 and 6 for JUND dimerized with ATF7, ATF2 and JUN. The second motif set, which is associated with JUND and FOSL2 or FOS, displays a single spacer at position 6. This spacer difference causes a bias in sequence alignment during the motif discovery process. It leads to a low informative PWM in the portion accounting for the TF partner when the motif is built with all JUND peaks. This example illustrates how *forkedTF* can systematically refine degenerated PWMs of a TF of interest built with a mixture of partners that require distinct spacer sizes for DNA binding. Additionally, when the co-factors are ranked based on the p-adjusted values (q-values) resulting from the enrichment test, the order remains the same for these five co-binding partners.

Finally, in this example, *forkedTF* mapped the methylation levels computed from WGBS data on the binding motif of JUND and its co-factors, shown as bar plots on the corresponding cytosines. We can observe that all sequences have low DNA methylation levels (Figure 2B).

### FPWM models co-factor binding sites appropriately and improves transcription factor binding prediction

We used transcription factor binding predictions to test FPWM’s ability to model JUND and its co-factors binding. After generating the FPWM for JUND and its partners in HepG2, we exported the results into the TRANSFAC format for the JUND+ATF2 and the JUND+FOSL2 motifs. These motifs were chosen because they represent the two distinctive sequences JUND can bind in HepG2 (double spacer and single spacer, respectively) (Figure 3). We extracted the genomic sequences corresponding to overlapping JUND and ATF2 peaks, as well as overlapping JUND and FOSL2 peaks. First, we evaluated the FPWMs of JUND, JUND+ATF2 and JUND+FOSL2 through cross-validation of their score distributions in each overlapping peak set (Figure 4B). More precisely, we compared the scores of an empirical distribution (red curve) against a permuted distribution (blue curve) and a theoretical (gray curve) across a dynamic range of weighted thresholds. Notice that the JUND+ATF2 motif empirical distribution curve (red) has the largest difference from the permuted curve (blue) in the JUND peaks overlapping ATF2 peaks dataset (JUNDxATF2 peak), implying that the JUND+ATF2 motif best model binding in JUND+ATF2 DNA sequences. Similar results can be observed for the JUND+FOSL2 motif in JUND peaks overlapping with FOSL2 peaks. Next, we evaluated the performance of co-factor motifs to predict binding sites. We observed that the JUND+ATF2 motif can predict a high number of binding sites in the DNA sequence of JUND peaks overlapping with ATF2 peaks (Figure 4C). This JUND+ATF2 binding strikingly contrasts with the low number of predicted binding sites obtained with the JUND+FOSL2 motif. The JUND motif calculated from all of the ChIP-seq peaks in HepG2 has a moderate performance. The reverse was observed when predicting binding in JUND peaks overlapping with FOSL2 peaks, where the JUND+FOSL2 has the best performance, and JUND+ATF2 has the worst prediction power (Figure 4D). In addition to obtaining a higher number of predicted binding sites, a better prediction was also achieved. This can be attributed to the use of a specific co-factor motif in their respective overlapping peak sequences, which is translated into more significant predictions, as can be observed in the p-values of (Figure 4E and 4F), particularly at the centers of the peaks, where the binding site is more likely to be present. Similar results were observed in other datasets, whereby motifs built from FPWMs outperform those constructed from PWMs in predicting binding sites (Supplementary Figure 4). Altogether, our results show that FPWM better model TF binding affinities of binding partners than PWMs.

**Figure 3.**
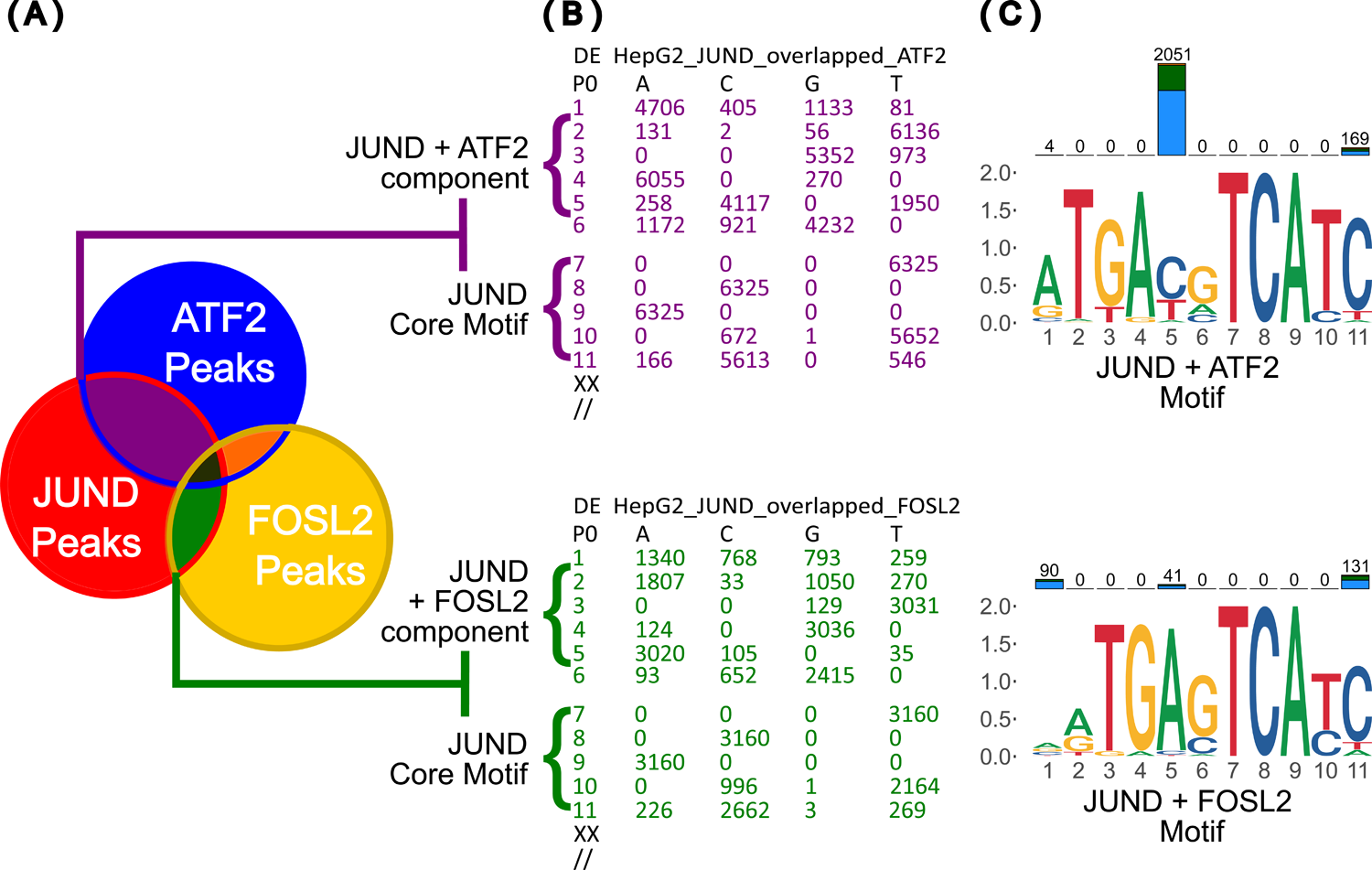
Building an FPWM object. **(A)** FPWM objects are generated by analyzing co-binding events across a TF of interest (here CEBPB) and its partners for dimerization (here ATF4 and CEBPD). By default, PWMs for partners are generated from sequences that overlap with the main TFBS. In contrast, PWM for the TF of interest, which is supposed to be conserved across all the binding partners, is built from combining sequences that overlap ChIP-seq binding sites from all co-binding partners. **(B)** TFregulomeR exports PWMs in TRANSFAC format for intersected peak regions. Here, the core motif and variable spacer are clearly present in the matrix. **(C)** TFregulomeR can also generate motifs for the above PWMs. The motifs based on intersected matrices have less noise in the classically degenerated section of the motif. The motif also captures the dinucleotide spacers’ propensity to form CpGs.

**Figure 4.**
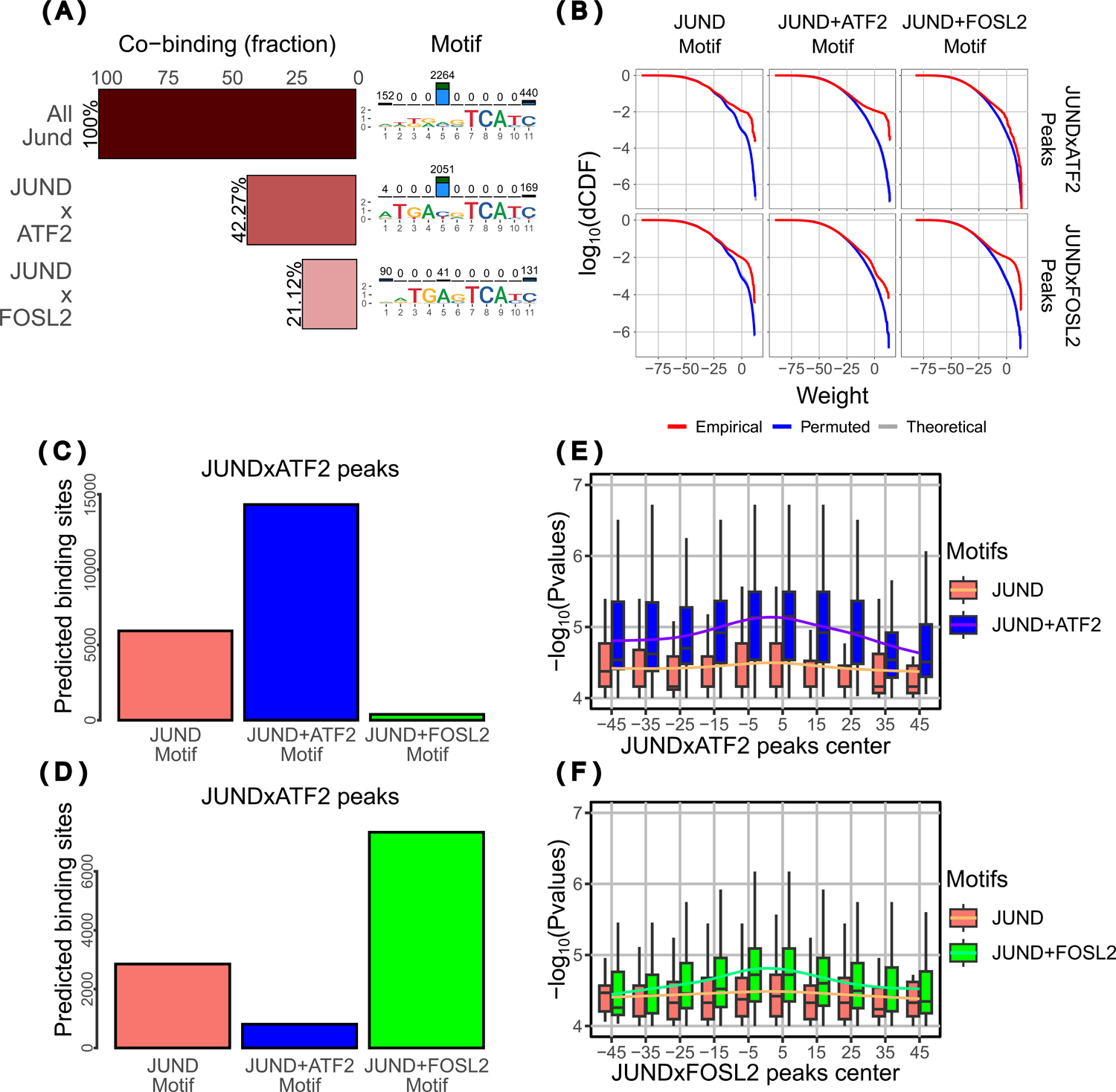
Comparative enrichment analysis for selected JUND heterodimer motifs. **(A)** *MiniCofatorReport* for two binding partners that yield distinctive spacer motifs. When JUND is bound with ATF2, the TGA and TCA motif halves are separated by a variable two-nucleotide spacer. When JUND is bound with FOSL2, the motif halves are separated by a single nucleotide spacer. **(B)** Score distributions were obtained when the JUNDxATF2 and JUNDxFOSL2 peaks with JunD, JUND+ATF2 and JUND+FOSL2 count matrices, respectively. The higher the separation between the empirical curve (red), the theoretical curve (inferred from dinucleotide frequencies, in grey) and the negative control curve (using randomly permuted matrices, in blue), the more specific the particular matrix is to a set of peaks. **(C)** Using the three different matrices, the number of predicted binding sites in the JUNDxATF2 peaks. **(D)** Using the three different matrices, the number of predicted binding sites in the JUNDxFOSL2 peaks. **(E)** P-value distribution of the matrix-scan results using JUND and JUND+ATF2 matrices around the center of JUNDxATF2 peaks. **(F)** P-value distribution of the matrix-scan results using JUND and JUND+FOSL2 matrices around the center of JUNDxFOSL2 peaks.

### *forkedTF* allows the detection of methylated DNA binding preferences in a co-factor specific way

Just like how a motif recognized by a TF may change depending on its co-binding partner, the ability of a TF to bind to methylated DNA may be modulated by its binding with a co-factor. We witness this modulation in the subset of DNA sequences bound by CEBPB and CEBPD complexes in K562 cells (Figure 5). We can observe sequence conservation across all partners in the first six bases. However, although position six retains cytosine, hypermethylation is present in the binding motif when CEBPB and CEBPD are bound together. This methylation signature and the emergence of a CpG dinucleotide in the motif appear only when building a FPWM focusing on the signal corresponding to CEBPB+CEBPD co-binding.

**Figure 5.**
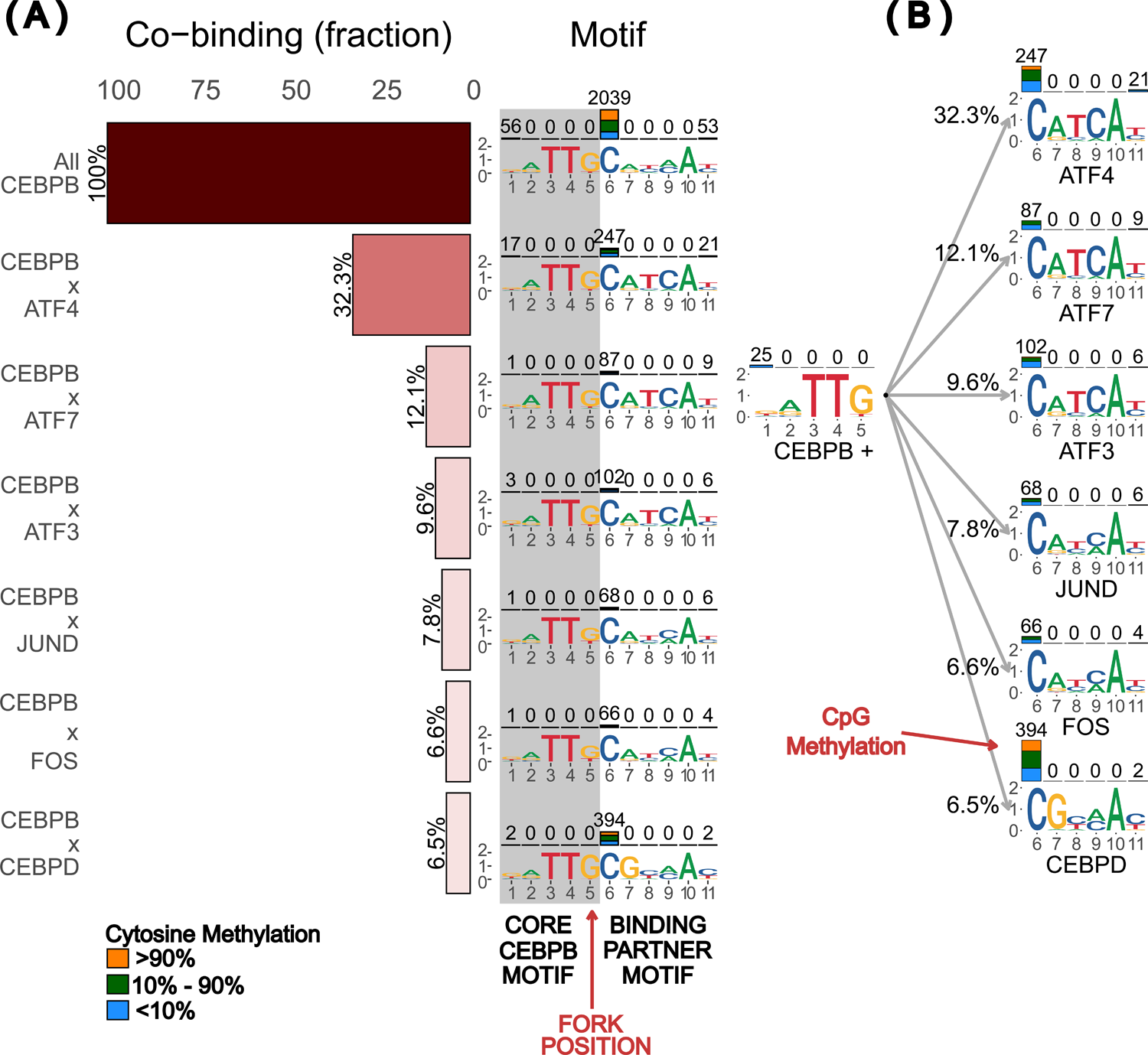
forkedTF of CEBPB in K562. **(A)** *miniCofactorReport* of CEBPB in K562 cells with its top six co-binding partners. **(B)** forkedLogo built with CEBPB and its main partners. We can observe that there is sequence conservation across all partners in the first six bases. However, despite the sequence conservation for the cytosine in position six, hypermethylation is present in the binding motif when CEBPB and CEBPD are bound together. This methylation signature and the emergence of a CpG dinucleotide in the motif cannot be observed without the binding partner segregation.

### Integration of *forkedTF* results within *MethMotif*

The results obtained with *forkedTF* for all 97 Human leucine zipper TFs are now integrated into the *MethMotif 2024* database. This extension includes the implementation of pre-computed FPWM, leveraging the advancements made by *forkedTF* in modeling the interactions between transcription factors and their co-factors. This information is available on *MethMotif* enhanced TF cards, and includes several new elements designed to provide a more detailed and nuanced understanding of transcription factor binding dynamics in the context of DNA methylation. Firstly, the incorporation of FPWM enables the modeling of transcription factor binding specificities when interacting with different co-factors. This addition allows for more accurate predictions of how TFs bind to DNA, reflecting the complex interplay between TFs binding partners and DNA methylation.

Within each motif card, users will find F-Logos, which are graphical representations illustrating both the core and variant motifs that TFs recognize under different conditions. Additionally, the motif cards provide a standard TRANSFAC file, which includes comprehensive details on the core motif as well as its leaf motifs, offering a detailed view of the TF binding preferences. Moreover, the FPWM TRANSFAC format is included, providing the specific data format needed to apply FPWM in various bioinformatic analyses (Supplementary Figure 5).

## DISCUSSION

In this manuscript, we highlight the crucial role of modeling partner-specific TFBS and showcase how *forkedTF* can identify the various DNA binding motifs that a TF can recognize in the presence of different binding partners. This approach generates accurate models of TF binding affinity, significantly enhancing the bioinformatic prediction of their binding sites. Additionally, *forkedTF* allows users to investigate the binding partners for all the available ChIP-seq datasets in a particular cell type, which has the possibility of uncovering novel TF-TF interactions. Moreover, we demonstrate that integrating an additional layer of epigenetic information, specifically DNA methylation, enables the identification of methyl-specific binding dimers. For instance, we show that the transcription factor CEBPB binds to a methylated DNA motif only when paired with the partner CEBPD.

While *forkedTF* relies on TFregulomeR to characterize co-factors, it can be easily adapted to create forked-PWM based on other approaches, such as PWM clustering, dyad motif discovery (12), or co-factor motif discovery. This flexibility is due to *forkedTF*’s ability to produce outputs in standard TRANSFAC format.

The effectiveness of this method relies entirely on the availability of public ChIP-seq datasets. The five cell lines documented with the highest number of TF ChIP-seq data sets available are HepG2 with 651, K562 with 628, A549 with 243 and GM12878 with 188 ChIP-seq datasets. Hence, we are still facing a very low coverage of the binding of the 1, 600 TF in human tissues (13). Nonetheless, even though our bioinformatic predictions are based on incomplete data, we were able to address a recurrent issue of sequence degeneration in PWM. As new datasets are continuously deposited in these repositories, predictions made by *forkedTF* will progressively improve, thereby enhancing our understanding of TF cooperativity.

Our method relies on co-binding, which means observing two TFs binding at the same genomic loci on many occasions across the genome. While there is evidence of heterodimerization for many of the TF pairs considered above, e.g. JUN+ATF2, JUND+FOS and CEBPB+CEBPD, this might not be true for other dimer pairs. TF binding close to one another in the same region to regulate the same genes may display similar results. Hence, users should be cautious not to interpret this as a confirmation of physical interaction between the two TFs.

It is now well-established that DNA methylation plays a critical role in orchestrating TF binding (14). Integrating DNA methylation patterns with DNA binding motifs has become a crucial step in enhancing our understanding of the dynamic interactions between TFs and their binding sites. This integration provides deeper insights into the regulatory mechanisms of chromatin structure and gene expression. While our *forkedTF-MethMotif* framework effectively illustrates the influence of DNA methylation on TF binding, tools that comprehensively scan and predict binding sites by incorporating this epigenetic information are still in the early stages of development (15, 16). We anticipate expanding our framework to integrate these aspects into a more robust model for transcription factor binding prediction in the future.

## DATA AVAILABILITY

The forkedTF package (encoded in R) and a user manual, along with all the scripts used in the reported analyses, are available on GitHub (https://github.com/benoukraflab/forkedTF).

## ACKNOWLEDGEMENTS

The authors thank Zoha Rabie for her proof-reading as well as Michael Woods and Jacques van Helden for advice and discussion. This research was enabled in part by support provided by ACENET (www.ace-net.ca) and Digital Research Alliance of Canada (ccdb.alliancecan.ca), as well as the Centre for Analytics, Informatics and Research (CAIR) at Memorial University.

## Authors contributions

M.D., R.T.M., A.G.K. and T.B. designed the program; M.D., R.T.M. and A.G.K. programmed the library and performed all analysis, with additional contributions from Q.X.X.L. and D.T.; W.S. implemented the q-value function; M.D. generated the Forked-Sequence Logos for Methmotif.org; all authors were involved in results interpretation; M.D., R.T.M., D.T., and T.B. wrote the manuscript; T.B. directed the project.

## REFERENCES

1. Stormo, G.D. (2000) DNA binding sites: representation and discovery. Bioinformatics, 16, 16–23.

2. Schneider, T.D. and Stephens, R.M. (1990) Sequence logos: a new way to display consensus sequences. Nucleic Acids Res., 18, 6097–6100.

3. Klein, D.C. and Hainer, S.J. (2020) Genomic methods in profiling DNA accessibility and factor localization. Chromosome Res., 28, 69–85.

4. Lin, Q.X.X., Thieffry, D., Jha, S. and Benoukraf, T. (2020) TFregulomeR reveals transcription factors’ context-specific features and functions. Nucleic Acids Res., 48, e10–e10.

5. Xuan Lin, Q.X., Sian, S., An, O., Thieffry, D., Jha, S. and Benoukraf, T. (2019) MethMotif: an integrative cell specific database of transcription factor binding motifs coupled with DNA methylation profiles. Nucleic Acids Res., 47, D145– D154.

6. Dyer, M., Lin, Q.X.X., Shapoval, S., Thieffry, D. and Benoukraf, T. (2024) MethMotif.Org 2024: a database integrating context-specific transcription factor-binding motifs with DNA methylation patterns. Nucleic Acids Res., 52, D222–D228.

7. Yevshin, I., Sharipov, R., Valeev, T., Kel, A. and Kolpakov, F. (2017) GTRD: a database of transcription factor binding sites identified by ChIP-seq experiments. Nucleic Acids Res., 45, D61–D67.

8. Lawrence, M., Huber, W., Pagès, H., Aboyoun, P., Carlson, M., Gentleman, R., Morgan, M.T. and Carey, V.J. (2013) Software for Computing and Annotating Genomic Ranges. PLoS Comput. Biol., 9, e1003118.

9. Machanick, P. and Bailey, T.L. (2011) MEME-ChIP: motif analysis of large DNA datasets. Bioinformatics, 27, 1696–1697.

10. Santana-Garcia, W., Castro-Mondragon, J.A., Padilla-Gálvez, M., Nguyen, N.T.T., Elizondo-Salas, A., Ksouri, N., Gerbes, F., Thieffry, D., Vincens, P., Contreras-Moreira, B., et al. (2022) RSAT 2022: regulatory sequence analysis tools. Nucleic Acids Res., 50, W670–W676.

11. Medina-Rivera, A., Abreu-Goodger, C., Thomas-Chollier, M., Salgado, H., Collado-Vides, J. and Van Helden, J. (2011) Theoretical and empirical quality assessment of transcription factor-binding motifs. Nucleic Acids Res., 39, 808–824.

12. van Helden, J., Rios, A.F. and Collado-Vides, J. (2000) Discovering regulatory elements in non-coding sequences by analysis of spaced dyads. Nucleic Acids Res., 28, 1808–1818.

13. Davis, C.A., Hitz, B.C., Sloan, C.A., Chan, E.T., Davidson, J.M., Gabdank, I., Hilton, J.A., Jain, K., Baymuradov, U.K., Narayanan, A.K., et al. (2018) The Encyclopedia of DNA elements (ENCODE): data portal update. Nucleic Acids Res., 46, D794–D801.

14. Tirado-Magallanes, R., Rebbani, K., Lim, R., Pradhan, S. and Benoukraf, T. (2017) Whole genome DNA methylation: beyond genes silencing. Oncotarget, 8, 5629–5637.

15. Liu, M.-L., Su, W., Wang, J.-S., Yang, Y.-H., Yang, H. and Lin, H. (2020) Predicting Preference of Transcription Factors for Methylated DNA Using Sequence Information. Mol. Ther. Nucleic Acids, 22, 1043–1050.

16. Nishizaki, S.S. and Boyle, A.P. (2022) SEMplMe: a tool for integrating DNA methylation effects in transcription factor binding affinity predictions. BMC Bioinformatics, 23, 317.

