## Supplementary Figures for "Representing Transcription Factor Dimer Binding Sites Using Forked-Position Weight Matrices and Forked-Sequence Logos"

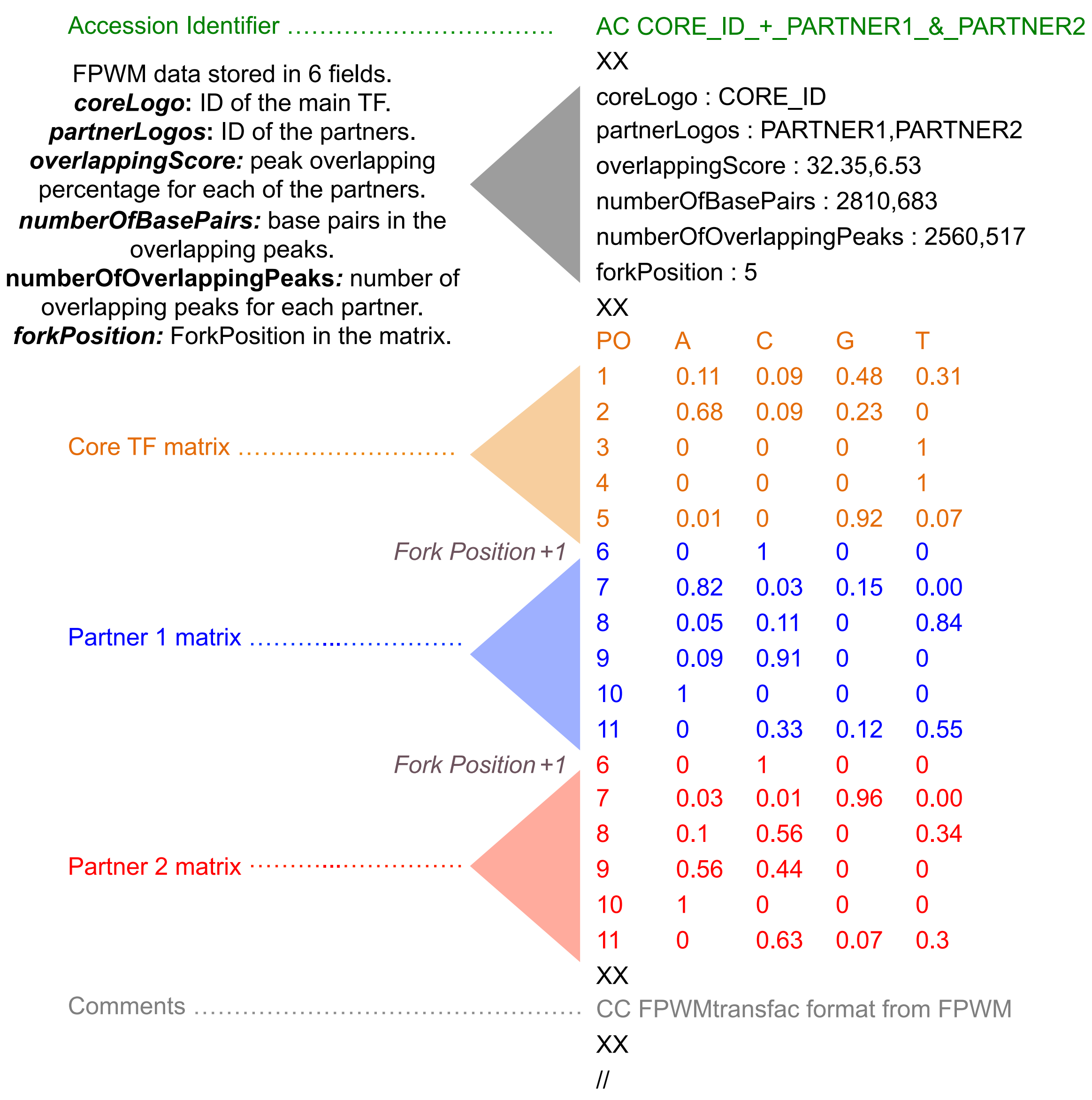


**Supplementary Figure 1. The FPWM format.**
This format is based on the TRANSFAC PWM format. It includes a metadata section that lists the IDs of the core TF and its partners, followed by information on its construction (overlapping percentage between the main TF and partner, number of base pairs and number of peaks in the overlap), and the fork position. The following section records the PWM, starting with the core TF matrix up to the fork position and followed by the matrices of the binding partners. All partner matrix positions starts from the fork position + 1. The file ends with an optional section for comments.

**(A)** **Count Matrix**

AC MM1_K562_CEBPB_+_MM1_K562_ATF4_&_MM1_K562_CEBPD
XX
parentLogo : MM1_HSA_K562_CEBPB
leafLogos : MM1_HSA_K562_ATF4,MM1_HSA_K562_CEBPD
overlappingScore : 32.3477381854941,6.53272681324236
numberOfBasePairs : 2810,683
numberOfOverlappingPeaks : 2560,517
forkPosition : 5
XX

| | P0 | A | C | G | T |
|---|---|---|---|---|---|
| CEBPB Core Motif | 1 | 349 | 274 | 1427 | 915 |
| | 2 | 2009 | 270 | 686 | 0 |
| | 3 | 0 | 0 | 0 | 2965 |
| | 4 | 0 | 0 | 0 | 2965 |
| | 5 | 29 | 0 | 2714 | 222 |
| + ATF4 Motif | 6 | 0 | 2560 | 0 | 0 |
| | 7 | 2106 | 68 | 374 | 12 |
| | 8 | 130 | 269 | 0 | 2161 |
| | 9 | 231 | 2329 | 0 | 0 |
| | 10 | 2560 | 0 | 0 | 0 |
| | 11 | 0 | 843 | 306 | 1411 |
| + CEBPD Motif | 6 | 0 | 517 | 0 | 0 |
| | 7 | 15 | 4 | 497 | 1 |
| | 8 | 50 | 288 | 0 | 179 |
| | 9 | 287 | 230 | 0 | 0 |
| | 10 | 517 | 0 | 0 | 0 |
| | 11 | 0 | 327 | 37 | 153 |

XX
CC transfacFormat: FPWMtransfac format from FPWM
CC matrixFormat: Count matrix
XX
//

**(B)** **Probability Matrix**

AC MM1_K562_CEBPB_+_MM1_K562_ATF4_&_MM1_K562_CEBPD
XX
parentLogo : MM1_HSA_K562_CEBPB
leafLogos : MM1_HSA_K562_ATF4,MM1_HSA_K562_CEBPD
overlappingScore : 32.3477381854941,6.53272681324236
numberOfBasePairs : 2810,683
numberOfOverlappingPeaks : 2560,517
forkPosition : 5
XX

| | P0 | A | C | G | T |
|---|---|---|---|---|---|
| CEBPB Core Motif | 1 | 0.11 | 0.09 | 0.48 | 0.31 |
| | 2 | 0.68 | 0.09 | 0.23 | 0 |
| | 3 | 0 | 0 | 0 | 1 |
| | 4 | 0 | 0 | 0 | 1 |
| | 5 | 0.01 | 0 | 0.92 | 0.07 |
| + ATF4 Motif | 6 | 0 | 1 | 0 | 0 |
| | 7 | 0.82 | 0.03 | 0.15 | 0.00 |
| | 8 | 0.05 | 0.11 | 0 | 0.84 |
| | 9 | 0.09 | 0.91 | 0 | 0 |
| | 10 | 1 | 0 | 0 | 0 |
| | 11 | 0 | 0.33 | 0.12 | 0.55 |
| + CEBPD Motif | 6 | 0 | 1 | 0 | 0 |
| | 7 | 0.03 | 0.01 | 0.96 | 0.00 |
| | 8 | 0.1 | 0.56 | 0 | 0.34 |
| | 9 | 0.56 | 0.44 | 0 | 0 |
| | 10 | 1 | 0 | 0 | 0 |
| | 11 | 0 | 0.63 | 0.07 | 0.3 |

XX
CC transfacFormat: FPWMtransfac format from FPWM
CC matrixFormat: Probability matrix
XX
//

**(C)** **Scale Count Matrix**

AC MM1_K562_CEBPB_+_MM1_K562_ATF4_&_MM1_K562_CEBPD
XX
parentLogo : MM1_HSA_K562_CEBPB
leafLogos : MM1_HSA_K562_ATF4,MM1_HSA_K562_CEBPD
overlappingScore : 32.3477381854941,6.53272681324236
numberOfBasePairs : 2810,683
numberOfOverlappingPeaks : 2560,517
forkPosition : 5
XX

| | P0 | A | C | G | T |
|---|---|---|---|---|---|
| CEBPB Core Motif | 1 | 349 | 274 | 1427 | 915 |
| | 2 | 2009 | 270 | 686 | 0 |
| | 3 | 0 | 0 | 0 | 2965 |
| | 4 | 0 | 0 | 0 | 2965 |
| | 5 | 29 | 0 | 2714 | 222 |
| + ATF4 Motif | 6 | 0 | 2965 | 0 | 0 |
| | 7 | 2439 | 79 | 433 | 14 |
| | 8 | 151 | 312 | 0 | 2502 |
| | 9 | 268 | 2697 | 0 | 0 |
| | 10 | 2965 | 0 | 0 | 0 |
| | 11 | 0 | 976 | 354 | 1635 |
| + CEBPD Motif | 6 | 0 | 2965 | 0 | 0 |
| | 7 | 86 | 23 | 2850 | 6 |
| | 8 | 287 | 1651 | 0 | 1027 |
| | 9 | 1646 | 1319 | 0 | 0 |
| | 10 | 2965 | 0 | 0 | 0 |
| | 11 | 0 | 1876 | 212 | 877 |

XX
CC transfacFormat: FPWMtransfac format from FPWM
CC matrixFormat: Scale Count matrix
XX
//

**Supplementary Figure 2. FPWM output formats.**
Because standard TRANSFAC matrices are derived from a set of DNA sequences of the same length, the sum of all the nucleotide counts remains the same for every row. However, in FPWMs, binding sequences are split between the main TFBS and TFBS belonging to co-factors. Therefore, here, only the sum of co-factors binding sequences equals the sum of binding sequences of the main TF. To allow FPWM to be compatible with the majority of PWM-scan tools, we implemented three different matrix formats: (A) Count matrix, which records raw counts from the intersection of peaks. Therefore, the sum of elements in a row can differ across rows. (B) Probability matrix, where counts in rows are divided by the sum of the elements in these rows. Hence, the sum of elements in rows always equals 1. (C) Scaled Count matrix, where counts are scaled concerning the highest value among the sum of elements for each row.

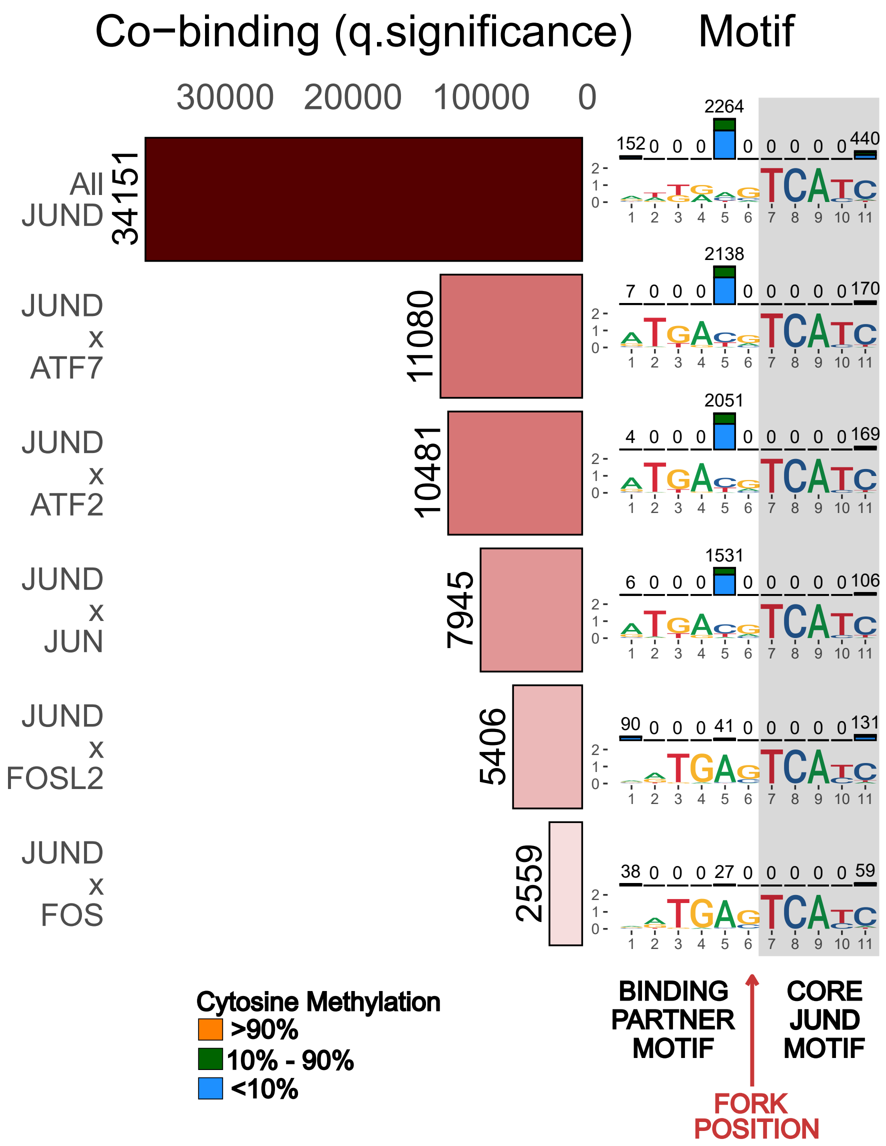


**Supplementary Figure 3. forkedTF q.significance analysis of JUND binding profile in HepG2 cell line.**

*miniCofactorReport* displays the results of ranking the binding partners based on the p-adjusted values (q-values) from an enrichment test. In the present bar chart, the q.significance is represented by -log10(q-value).

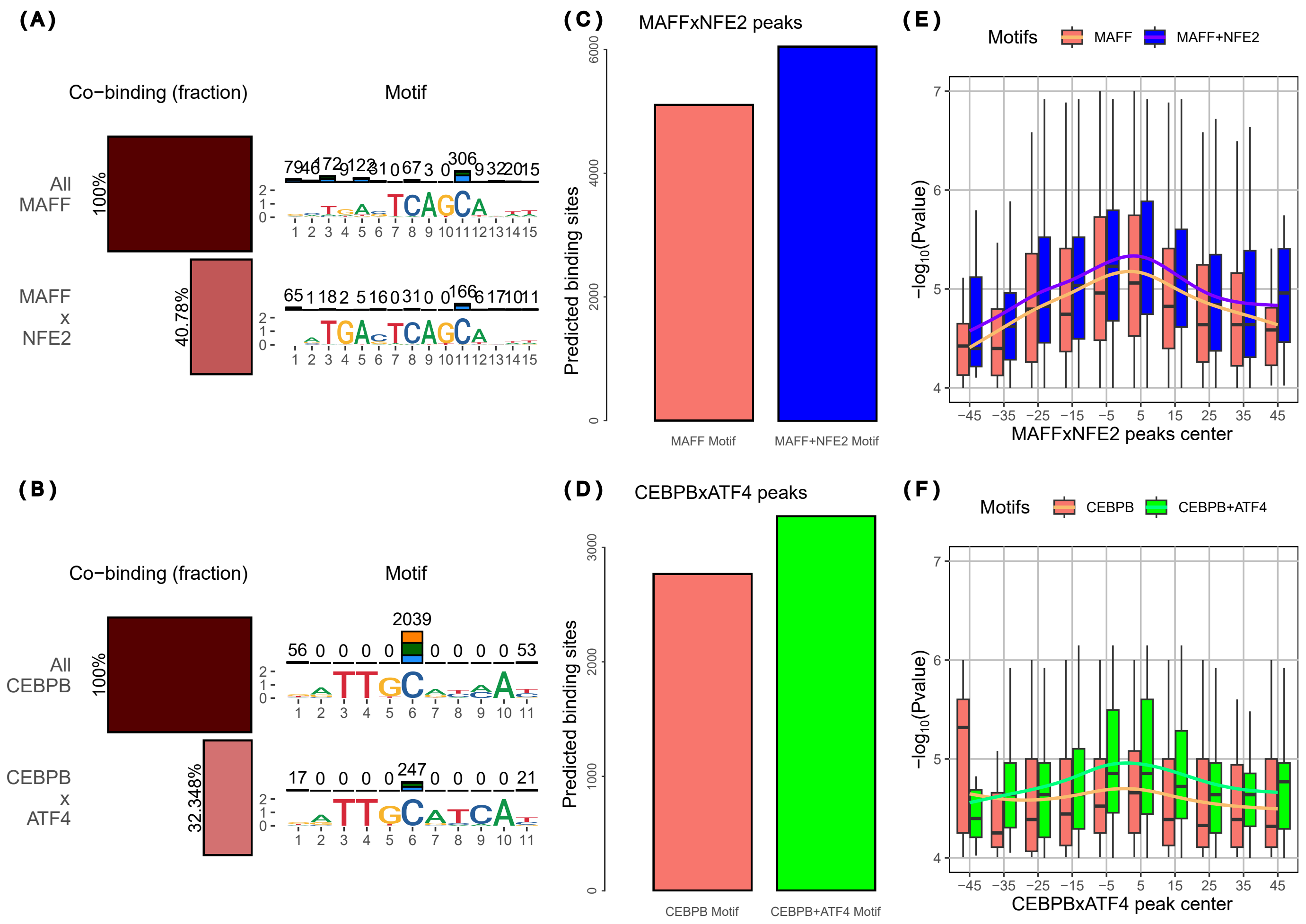


**Supplementary Figure 4. Examples of forkedTF matrix scans.**
(A) miniCofactorReport for MAFF and its binding partner NFE2 in the K562 cell line. (B) minicofactorReport CEBPB and its partner ATF4 in K562. (C) The number of predicted binding sites in the MAFF peaks that overlap with NFE2 peaks, using the matrix of all MAFF peaks and MAFFxNFE2 peaks. (D) The number of predicted binding sites in the CEBPB peaks that overlap with ATF4 peaks, using the matrix of all CEBPB peaks and CEBPBxATF4 peaks. (E) P-value distribution of the matrix-scan results using MAFF and MAFF+NFE2 motif around the peak center of MAFFxNFE2 peaks. (F) P-value distribution of the matrix-scan results using CEBPB and CEBPB+ATF4 motif around the peak center of CEBPBxATF4 peaks.

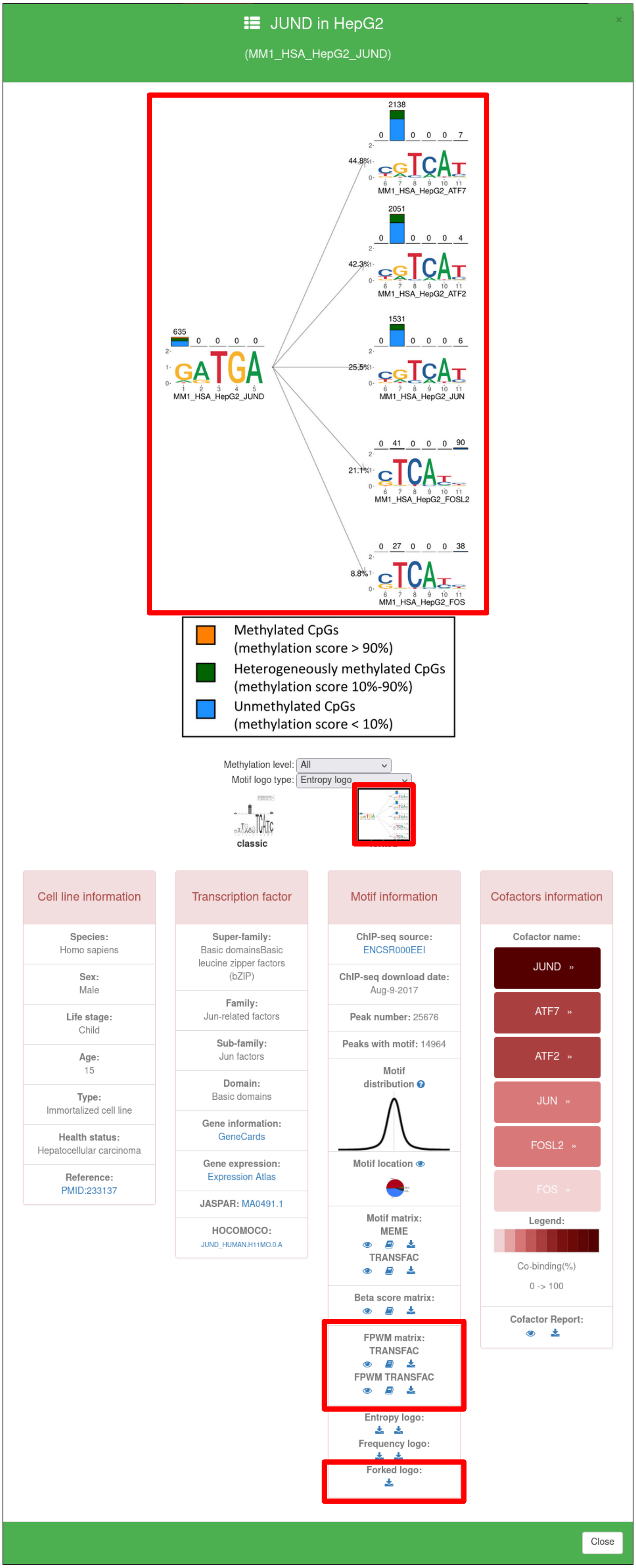


**Supplementary Figure 5. Implementation of the results of forkedTF analyses in MethMotif 2024.**
The screenshot of MethMotif highlights the new motif card within the MethMotif database. The main logo display can be toggled between the classic logo and the new F-Logo (upper two red boxes). Under Motif Information (bottom, middle-right), we further provide the corresponding motif in classic TRANSFAC and FPWM TRANSFAC formats (red box), as well as various graphical logo representations, including the new F-Logo representation (bottom red box). All these updates are currently available for the transcription factor dimers belonging to the bZIP family.
